# Inhibition of macrophage neuraminidase 1 protects against immune thrombocytopenia by limiting platelet clearance

**DOI:** 10.64898/2026.09.03.748973

**Authors:** Antoine Caillon, Lorena Carvelli, Xuefang Pan, Diogo Poroca, Silvia Neri, Elisa G. Carvajal, Aymeric Fabié, Manon Saby, Yves Pastore, Renée Bazin, Thomas Pincez, Christopher W. Cairo, Alexey V. Pshezhetsky

**Author notes:** Corresponding author: Alexey V. Pshezhetsky, CHU Sainte-Justine Research Center, 3175 Chemin de la Côte-Ste-Catherine, Montréal (QC) H3T 1C5, Canada. /2736.

## Abstract

Immune thrombocytopenia purpura (ITP) is an autoimmune disorder characterized by a reduction in circulating platelet levels, primarily due to generation of autoantibodies to platelet surface antigens followed by their spleen macrophage-mediated clearance. Emerging evidence implicates neuraminidase (sialidase) enzymes including neuraminidase 1 (NEU1) in platelet clearance and ITP severity; however, the underlying cellular mechanisms remain unknown. Using tissue-specific NEU1 knockout mouse models, we studied the contribution of platelet and macrophage NEU1 to ITP pathogenesis and evaluated whether pharmacological inhibition of NEU1 could preserve platelet counts in a murine ITP model. Constitutive and macrophage-specific, but not platelet-specific, NEU1 knockout mice showed a protection against reduction of platelet counts in the passive ITP model suggesting that macrophage, but not platelet, NEU1 promotes platelet clearance. Genetic deletion or pharmacological blockade of macrophage NEU1 also reduced platelet phagocytosis by cultured macrophages in vitro. The selective NEU1 inhibitor CG33301 protected mice against anti-CD41a antibody-induced thrombocytopenia and showed a higher efficacy compared to pan neuraminidase inhibitor oseltamivir phosphate. Our results demonstrate that the macrophage pool of NEU1 plays a central role in platelet clearance by splenocytes during ITP by activating their phagocytosis and suggest that selective NEU1 inhibition may be a promising therapeutic strategy for this disease.

## Introduction

Immune thrombocytopenia purpura (ITP) is an autoimmune condition defined by low platelet counts due to peripheral destructions of circulating platelets in the absence of bone marrow-related abnormalities. ITP is a common cause for low platelet counts in childhood affecting between 4-8 per 100,000 children each year with a mean age at presentation of 5.7 years [1–3]. ITP is an acute condition when it occurs in children, and thrombocytopenia will often resolve within a couple of weeks to months. However, up to 25% may have prolonged thrombocytopenia beyond 12 months of presentation, children older than 10 years of age being at increased risk to have chronic thrombocytopenia lasting more than 12 months [3, 4]. Variability in natural history and response to therapy suggest that ITP comprises heterogeneous disorders arising through diverse mechanisms, all involving production of anti-platelet autoantibodies. Historically, ITP has been attributed to formation of autoantibodies directed against surface platelet glycoproteins, particularly GPIIb/IIIa and GPIb/IX, which opsonize platelets for Fcγ receptor–mediated clearance by macrophages in the spleen and liver [5]. However, more recent studies demonstrated that the pathophysiological mechanism underlying ITP might also involve the desialylation of surface platelet glycoproteins mediated by human neuraminidase 1 (NEU1), a lysosomal sialidase mainly active on the terminal sialic acids of glycoproteins (reviewed in [6]). Desialylated glycoproteins on the platelet surface expose terminal galactose residues, recognized by the Ashwell-Morell (asialoglycoprotein) receptor and Macrophage galactose lectin (MGL) receptor of hepatocytes and liver macrophages, leading to clearance of platelets[7–16]. Notably, CD8(+) T cells of ITP patients with positive cytotoxicity reveal abundant NEU1 levels on the platelet surface, platelet desialylation and platelet phagocytosis by hepatocytes [17]. NEU1 as well as the cytoplasmic neuraminidase 2 (NEU2) were also reported to play an essential role in the platelet adherence to von Willebrand Factor (VWF) via the platelet surface glycoprotein Ibα (GPIbα) [18]. Consequently, pharmacological neuraminidase inhibition has emerged as a potential therapeutic avenue for ITP. Specifically, oseltamivir phosphate (OP), an inhibitor of influenza virus neuraminidase clinically approved for treatment of influenza under the commercial name Tamiflu, was reported to show partial efficacy in murine and humanized murine ITP models as well as in some ITP patients, particularly in the cases involving production of anti-GPIbα antibodies [11, 17, 19–21]. However, the biochemical mechanism underlying these results needed further clarification, especially considering that OP shows very low inhibitory activity against human and mouse NEU1 (IC > 1000 μM [22–24]) and may not directly inhibit the enzyme under the conditions used in the above experiments. On the other hand, NEU1 specific inhibitors showed efficacy in inhibiting the desialylation of low density lipoproteins and their uptake by macrophages via Ashwell-Morell receptor *in vivo* [12].

Multiple studies, including those from our laboratories, demonstrated that beyond its canonical role in the lysosomal catabolism of sialoglycoproteins and oligosaccharides, NEU1 also cleaves sialic acid residues from the glycan chains of receptors at the plasma membrane, which may change their conformation, half-life on the cell surface and biological activity, and modulate receptor signaling, related to immunity, inflammation and cellular phagocytosis [12, 25–28] (reviewed in [29]). In particular, we have shown that NEU1 desialylates and activates the macrophage FcγR receptor responsible for phagocytosis of IgG-opsonized particles[30]. Spleen macrophages from NEU1-deficient mice showed increased sialylation of FcγR receptor and impaired phagocytosis of IgG-coated red blood cells, while treatment with the exogenous NEU1 reduced sialylation of FcγR and restored phagocytosis[30]. Thus, inhibiting macrophage NEU1 activity could potentially block the uptake of antibody-opsonized platelets by splenocytes through FcγR receptors.

In the current study, we demonstrate that genetic inactivation of macrophage, but not platelet, NEU1 reduces the rate of platelet depletion in the mouse passive ITP model and the uptake of opsonized thrombocytes by spleen macrophages in vitro. We also show that NEU1-specific inhibitors exhibit higher potency in rescuing platelet counts compared to OP, suggesting that pharmacological inhibition of the enzyme may become a promising therapeutic strategy for ITP.

## Methods

### Study approval

Approval for animal experimentation was granted by the Animal Care and Use Committee of the CHU Ste-Justine (approval numbers 2022-3452 and 2022-3453).

### Animals

Constitutive KO (*Neu1*^Δ*Ex3*^), phagocyte-specific conditional (*Neu1^Cx3cr1^*^Δ*Ex3*^) and the cathepsin A hypomorph galactosialidosis (*CathA^S109A-Neo^*) mouse strains were previously described[31, 32]. A platelet-specific conditional KO (*Neu1^Pf4^*^Δ*Ex3*^) mouse was generated by crossing *Neu1^loxPEx3^* strain with *Neu1* exon 3, flanked with the loxP sites [31], with the C57BL/6-*^Tg(Pf4-icre)Q3Rsko^*/J strain expressing Cre recombinase under control of the *Pf4* (platelet factor 4) promoter. To generate inducible Neu1 KO (*Neu1*^Δ*Ex3-ind*^) mice, *Neu1^loxPEx3^* strain was crossed with B6.Cg-*Ndor1^Tg(UBC-cre/ERT2)1Ejb^*/1J strain expressing the Cre-ERT2 fusion gene under the control of the human ubiquitin C promoter. Upon treatment with tamoxifen, *Neu1* exon 3 is deleted in all tissues of *Neu1*^Δ*Ex3-ind*^ mice.

### Isolation of mouse spleen macrophages, Kupffer cells, hepatocytes and platelets

Spleen macrophages were isolated by dissociating spleen cells followed by removal of non-adherent cells. Only the batches containing >85% of CD11b^+^ cells with viability >90% were used. Liver hepatocytes and Kupffer cells were isolated as described [33–36] (see supplementary materials for details). Platelets were isolated as described [11] from the whole blood, collected by terminal bleeding from the heart.

### Flow cytometry

Cells labeled with viability dyes and treated with rat anti-mouse CD16/CD32 Fc receptor block were incubated with specific antibodies (Table S1) in PBS with 5% of fetal bovine serum, fixed, washed and analyzed by flow cytometry using a BD LSRFortessa cell analyzer (BD Biosciences). Data analysis was performed using the FlowJo software (version 10.1). Gating strategies are shown in supplementary materials.

### Analysis of platelet *in vitro* uptake

Platelets (10^6^ cells/mL) were labeled by PKH26 dye, premixed with anti-mouse CD41 monoclonal antibody (5 μg/ml), added to 10^5^ of WT of cultured macrophages and incubated at 37°C or 4°C for 30 min. Harvested macrophages were analyzed by flow cytometry to quantify the percentage of PKH26-positive macrophages and the mean fluorescence intensity. In selected experiments, macrophages were pretreated for 24 h or 30 min with NEU1 inhibitors: oseltamivir phosphate (OP), CG25901, CG33301, CG33300, and C9-BA-DANA prepared as described [28]. Platelet uptake by primary mouse hepatocytes and Kupffer cells was studied using a similar protocol with 2.6 10^5^ platelets loaded on 10^5^ phagocytic cells and incubated for 30 min at 4 or 37 °C. The experimental design and gating controls are shown in Fig. S1, S5 and S6.

### Passive murine ITP model

ITP in mice was induced as described by Katsman et al. [37]. The mice were injected IP with anti-mouse CD41 (68 μg/kg BW on the 1st day, 102 μg/kg BW on the 2d day) or anti-mouse CD42 (2 mg/kg BW) monoclonal antibodies. The reticulated platelets were measured by flow cytometry. Neuraminidase inhibitors were administered 1 h before and 2 h after the injection of anti-CD41a antibodies (Fig. S2).

### Statistical analysis

Statistical analyses were performed using Prism GraphPad 9.3.0. software using t-test, one-way ANOVA or Nested ANOVA tests (normal distribution) or Mann-Whitney and Kruskal-Wallis tests. A *P*-value of 0.05 or less was considered significant.

## Results

### Basal platelet count and bleeding time is altered in mice with a constitutive NEU1 deficiency but not in conditional platelet or macrophage-specific *Neu1* knockouts

Using constitutive and conditional *Neu1* gene-targeted mouse models, we tested whether NEU1 deficiency in platelets or macrophages influences platelet homeostasis under steady-state conditions. We used previously described constitutive *Neu1* KO (*Neu1*^Δ*Ex3*^), gene-targeted (*Neu1^Geo^*), conditional KO *Neu1^Cx3cr1^*^Δ*Ex3*^ with NEU1 deficiency in phagocytic mononucleosis cells, including liver, spleen and lung macrophages [31], and the galactosialidosis *CathA^S190A-Neo^* (cathepsin A hypomorph) strains [32]. Since NEU1 depends on cathepsin A in expression of its enzymatic activity [38] the residual NEU1 activity in tissues of *CathA^S190A-Neo^* mice is reduced to ∼9% of normal levels [32]. This level of residual NEU1 activity protects mice from lysosomal storage, but it is not sufficient for desialylation of receptors on the plasma membrane [30]. The conditional platelet-specific *Neu1^Pf4^*^Δ*Ex3*^ KO strain was generated in the current work by crossing previously described *Neu1^loxPEx3^* strain [31] with the C57BL/6-*^Tg(Pf4-icre)Q3Rsko^*/J strain expressing the Cre recombinase under the control of the *Pf4* (platelet factor 4) gene promoter. Analysis of total acidic neuraminidase (NEU1, NEU3 and NEU4), neutral neuraminidase (NEU2), and NEU1 specific activities revealed a ∼30% reduction in total acidic neuraminidase activity and almost complete absence of NEU1 specific activity in the platelets of *Neu1^Pf4^*^Δ*Ex3*^ mice (Fig. S3A). Similar results were observed for the platelets of constitutive *Neu1*^Δ*Ex3*^ mice. In contrast, the total neuraminidase activity in the liver and the kidney as well as the total neuraminidase and NEU1 activities in the bone marrow were reduced only in constitutive *Neu1*^Δ*Ex3*^ mice but not in *Neu1^Pf4^*^Δ*Ex3*^ mice compared to WT controls. Together, these results suggest specific genetic depletion of NEU1 in the platelets of *Neu1^Pf4^*^Δ*Ex3*^ mice. They also indicate that NEU1 in these cells shows a relatively low abundance (<30% of total activity) compared to other neuraminidases. The results of the activity assays were further supported by immunofluorescence microscopy that showed equally reduced areas labeled with anti-NEU1 antibodies in CD41^+^ platelets from both *Neu1^Pf4^*^Δ*Ex3*^ and *Neu1*^Δ*Ex3*^ mice compared to WT controls (Fig. S3B).

We further studied whether platelets of *Neu1^Pf4^*^Δ*Ex3*^ mice showed altered surface sialylation profiles. For this the cells were incubated with fluorescently labeled *Sambucus nigra* agglutinin (SNA), that specifically recognizes sialic acid residues that are attached to terminal galactose or N-acetylgalactosamine (GalNAc) via an α2,6 linkage. Cells were also labeled with Peanut Agglutinin (PNA) lectin binding to the terminal β-galactose residues, but not those masked by the terminal sialic acid residues. Wheat Germ Agglutinin (WGA) highly specific for N-acetyl-D-glucosamine (GlcNAc) was used as a control. In separate experiments, the cells were pretreated with an excess (10 mU/ml) of pan-specific neuraminidase from *Clostridium perfringens* in the absence of presence of pan-specific neuraminidase inhibitor, 2,3-didehydro-2-deoxy-N-acetyl-neuraminic acid (DANA; 1 mmol/L final concentration). Platelets from *Neu1^Pf4^*^Δ*Ex3*^ mice showed labeling with SNA and PNA lectins similar to WT mice, while platelets from constitutive *Neu1* KO mice showed increased SNA and reduced PNA labeling consistent with their increased sialylation (Fig. S4). Specificity of SNA was confirmed by treating the cells with bacterial neuraminidase which drastically reduced the labeling in all cells. WGA labeling was similar for platelets from all mice and was not changed by neuraminidase treatment (Fig. S4). Flow cytometry results were corroborated by the lectin blotting which revealed a similar affinity of platelet proteins to biotinylated PNA and SNA for WT and *Neu1^Pf4^*^Δ*Ex3*^ mice (Fig. S4).

The basal platelet levels in platelet-specific *Neu1^Pf4^*^Δ*Ex3*^ or macrophage-specific *Neu1^Cx3cr1^*^Δ*Ex3*^ KO mice were similar to those in the WT mice of similar genetic background, age and sex. In contrast, the constitutive *Neu1*^Δ*Ex3*^ *Neu1* KO mice displayed a significantly reduced platelet counts compared to WT counterparts (373.9 ± 54.0 ×10^3^/µL in *Neu1*^Δ*Ex3*^ vs 876.2 ± 39.7 ×10^3^/µL in WT; *p*<0.0001) (Fig. 1A). These results were confirmed by the hematology data (Fig. 1B) which showed reduced platelet levels in *Neu1*^Δ*Ex3*^, *Neu1^Geo^* and *CathA^S190A-Neo^* mice with constitutive NEU1 deficiency. In contrast, platelet counts in macrophage-specific *Neu1^Cx3cr1^*^Δ*Ex3*^ strain and platelet-specific *Neu1^Pf4^*^Δ*Ex3*^ KO strains as well as in the control *Neu1^loxPEx3^*strain were normal. Platelet counts in the neuraminidase 3 (*Neu3^-/-^*) and neuraminidase 4 (*Neu4^-/-^*) KO mice, as well as in the mouse model of lysosomal storage disease, Mucopolysaccharidosis IIIC (*Hgsnat^Geo^*) [39] were also similar to those of the WT mice. Other white blood cell counts were similar in all strains with an exception of monocytes which showed an increase in *Neu1*^Δ*Ex3*^ and *Neu1^Geo^* strains with a complete absence of NEU1 activity in tissues. These mice mimic human lysosomal disease sialidosis and manifest with progressive systemic inflammation [31]. Bleeding time assay did not reveal differences between WT, *Neu1^Pf4^*^Δ*Ex3*^, or *Neu1^Cx3cr1^*^Δ*Ex3*^ animals, but the constitutive KO *Neu1*^Δ*Ex3*^ mice exhibited a markedly prolonged bleeding time compared to WT mice (∼155 s vs ∼70 s; *p*<0.001), consistent with a reduced platelet count (Fig. 1C). To confirm association between systemic NEU1 deficiency and low basal levels of platelets, we have measured platelet counts in the inducible *Neu1* KO model *Neu1*^Δ*Ex3-ind*^ before and after the injection of tamoxifen, which results in the deletion of the *loxP*-flanked exon 3 from the *Neu1* gene and the progressive loss of the NEU1 activity in all tissues. Before the tamoxifen treatment, platelet counts in the *Neu1*^Δ*Ex3-ind*^ mice were similar to WT and *Neu1^loxPEx3^* controls, but 1 week after the treatment the platelet levels started to decrease and 4 weeks after reached ∼400 ×10^3^/µL level, similar to Neu1 KO mice (Fig. 1D). The platelet counts in control WT and *Neu1^loxPEx3^* mice were not affected by tamoxifen treatment (Fig. 1C). Together, these results indicate that systemic loss of NEU1 affects platelet levels and corroborate the reports of thrombocytopenia and thrombocytopathy in human sialidosis patients [40, 41]. In contrast, NEU1 deficiency only in platelets or phagocytic monocytes does not alter platelet levels and blood coagulation process.

**Figure 1.**
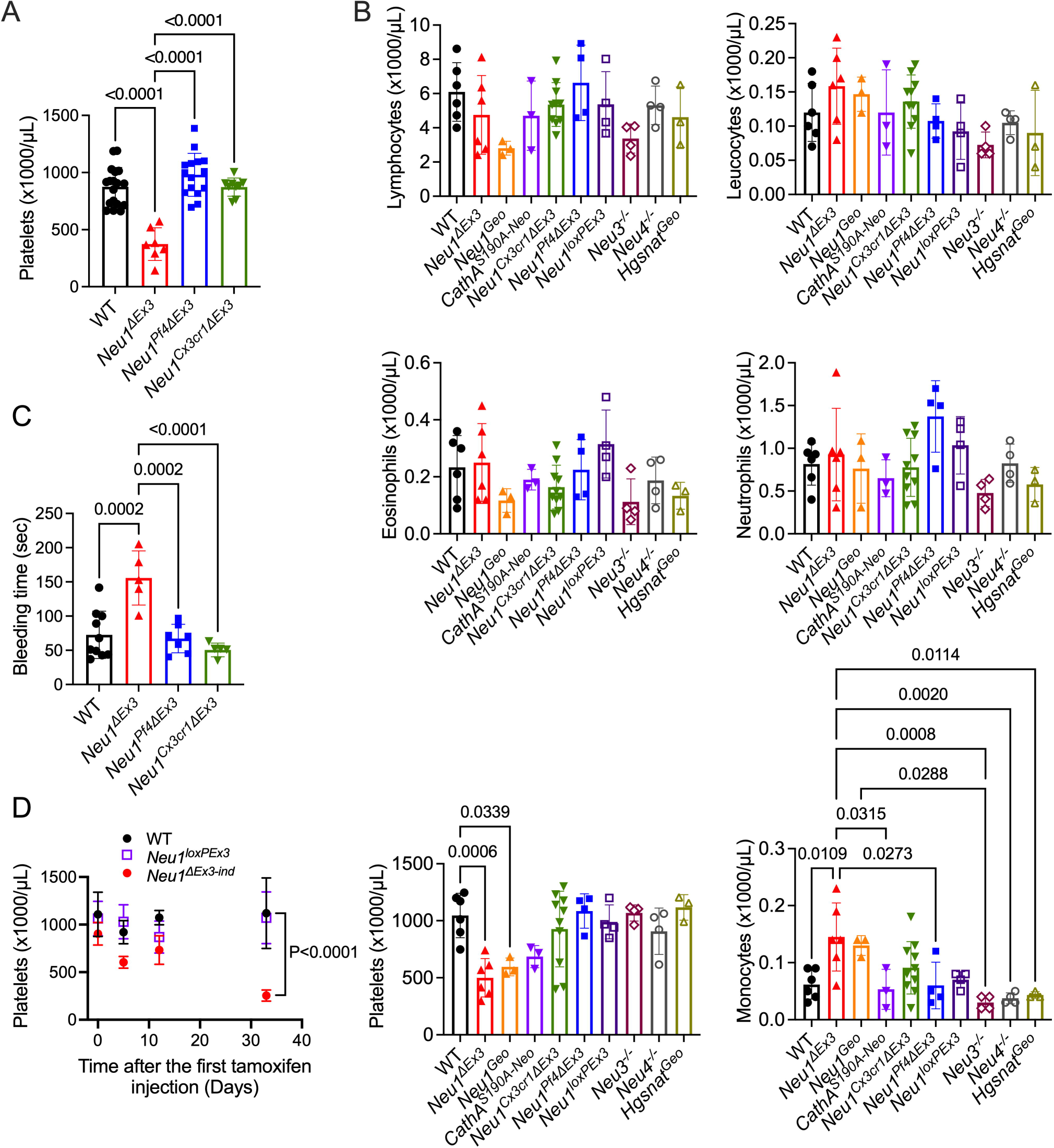
Constitutive NEU1 deficiency alters baseline platelet hemostasis. **(A)** Platelet counts in WT, constitutive *Neu1* KO (*Neu1*^Δ*Ex3*^), platelet KO (*Neu1^Pf4^*^Δ*Ex3*^), and macrophage KO (*Neu1^Cx3cr1^*^Δ*Ex3*^) mice. Constitutive but not macrophage or platelet-specific *Neu1* KO mice show reduced platelet counts. **(B)** Blood cell counts in WT and neuraminidase deficient mice. **(C)** Tail bleeding times in the same groups. Only constitutive *Neu1* KO mice show significantly prolonged bleeding time. **(D)** Induction of NEU1 deficiency in tamoxifen-inducible *Neu1* KO mice is associated with a reduction of platelet counts. WT and *Neu1^loxPEx3^* control mice injected with tamoxifen show stable levels of platelet counts. *P* values are calculated with one-way ANOVA (**A, B**) or two-way ANOVA (**C**) with Tukey and Dunnett’s multiple comparisons tests, respectively, individual results, means and SD for 5–17 mice/group are shown.

### Macrophage, but not platelet, NEU1 drives platelet clearance in the mouse model of ITP

To dissect the cellular role of NEU1 in immune thrombocytopenia (ITP), we used the passive mouse ITP model described by Katsman et al.[37]. Mice were receiving escalating daily doses (68 μg/kg on the first and 102 μg/kg on the second day) of anti-CD41a mouse monoclonal antiplatelet antibody (clone MWReg30) and the circulated platelets were measured before, 24 h, and 48 hours after the first antibody injection by flow cytometric analysis using the blood sample collected from the submandibular vein. Both constitutive *Neu1*^Δ*Ex3*^ and macrophage-specific *Neu1^Cx3cr1^*^Δ*Ex3*^ *Neu1* KO mice showed a significantly (*P*<0.0017 and <0.0012, respectively) slower decline rate of platelet counts compared to WT mice indicating that they were protected against thrombocytopenia (Fig. 2A). Galactosialidosis *CathA^S190A-Neo^* mice, which retain ∼9% of the residual NEU1 activity, displayed a similar phenotype (Fig. 2A). In contrast, platelet-specific *Neu1^Pf4^*^Δ*Ex3*^ KO showed the platelet depletion rate similar to that of the WT mice indicating that the removal of NEU1 only from thrombocytes offers no protection from platelet clearance (Fig. 2B). We, further, tested the rate of platelet decline induced by anti-CD42b (GPIbα) antibodies reported to be specific for the desialylated form of the protein. Mice were injected with 2 mg/kg BW of the anti-CD42b antibody and the concentration of platelets was measured in the blood collected before and 24 h after the injection. Like in the case of ITP induced by anti-CD41a antibody, we did not observe any difference in the platelet depletion rate between *Neu1^Pf4^*^Δ*Ex3*^ mice and their respective *Neu1^loxPEx3^*control (Fig. 2C). Together with the results of lectin analysis, this result suggests that the platelet NEU1 pool does not play a major role in desialylation of platelet surface proteins.

**Figure 2.**
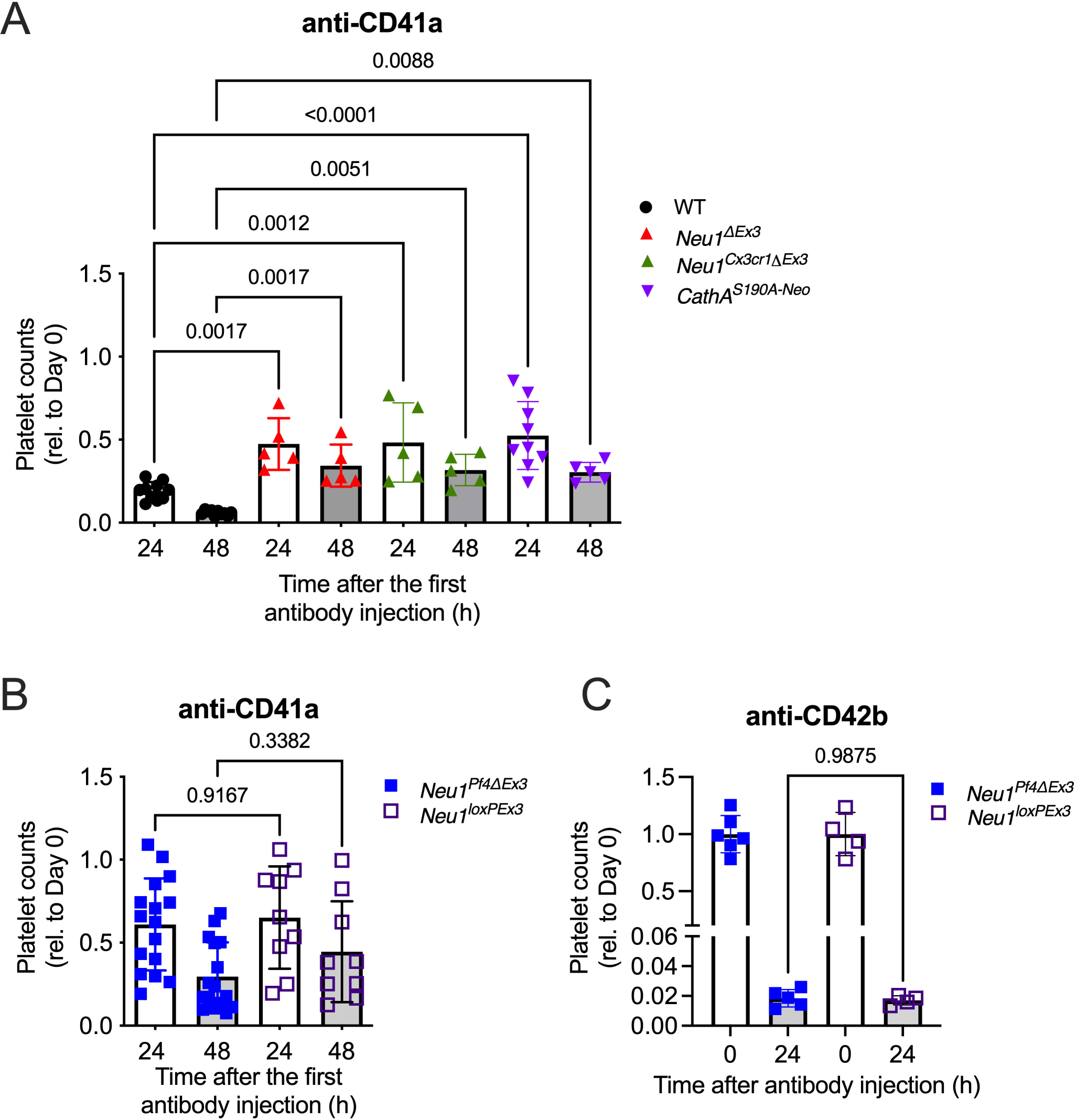
Constitutive or macrophage-specific NEU1 deficiency protects mice against ITP in the passive model. (A-B) Platelet counts in mice analysed before and after ITP induction with anti-CD41a antibody (68 μg/kg BW on day 1, 102 μg/kg on day 2). Constitutive KO (*Neu1*^Δ*Ex3*^), macrophage KO (*Neu1^Cx3cr1^*^Δ*Ex3*^) and galactosialidosis NEU1-deficient (*CathA^S190A-Neo^*) mice **(A)** but not platelet KO (*Neu1^Pf4^*^Δ*Ex3*^) mice **(B)** are partially protected against platelet immune depletion. **(C)** Platelet counts in mice analysed before and after ITP induction with anti-CD42b antibody (2 mg/kg BW on day 1). Platelet KO (*Neu1^Pf4^*^Δ*Ex3*^) mice show platelet depletion rate similar to that of control (*Neu1^loxPEx3^*) mice. Individual results, means ± SD (*n*=4-15) are shown. *P* values were calculated by one-way ANOVA with Šídák’s multiple comparisons test.

### Macrophage NEU1 promotes platelet uptake *in vitro*

To directly assess the rate of platelet uptake by macrophages, PKH26-labeled opsonized mouse platelets from WT or *Neu1^Pf4^*^Δ*Ex3*^ mice were added to cultured mouse splenic macrophages collected either from WT or *Neu1^Cx3cr1^*^Δ*Ex3*^ KO mice. After 30 min of incubation at 37 °C, macrophages were harvested, washed and analysed by flowcytometry to determine the fraction of cells containing the PKH26-labeled platelets and the relative intensity of labeling reflective of the number of platelets engulphed by or bound to each macrophage. Together, these parameters provide an integrated measure of platelet uptake, including both surface binding and internalization, which reflects functional platelet clearance. The macrophages from *Neu1^Cx3cr1^*^Δ*Ex3*^ mice showed a significantly reduced uptake (the percent of PKH26+ macrophages) compared to WT cells (Fig. 3A,B) regardless of the platelet source (WT or *Neu1^Pf4^*^Δ*Ex3*^ mice). The mean fluorescence intensity per cell was also reduced in the macrophages lacking NEU1, indicating that fewer platelets were ingested per macrophage (Fig. 3A,B). In contrast, similar uptake rate was detected for WT and NEU1-deficient platelets, harvested from the *Neu1^Pf4^*^Δ*Ex3*^ mice (Fig. 3A,B).

**Figure 3.**
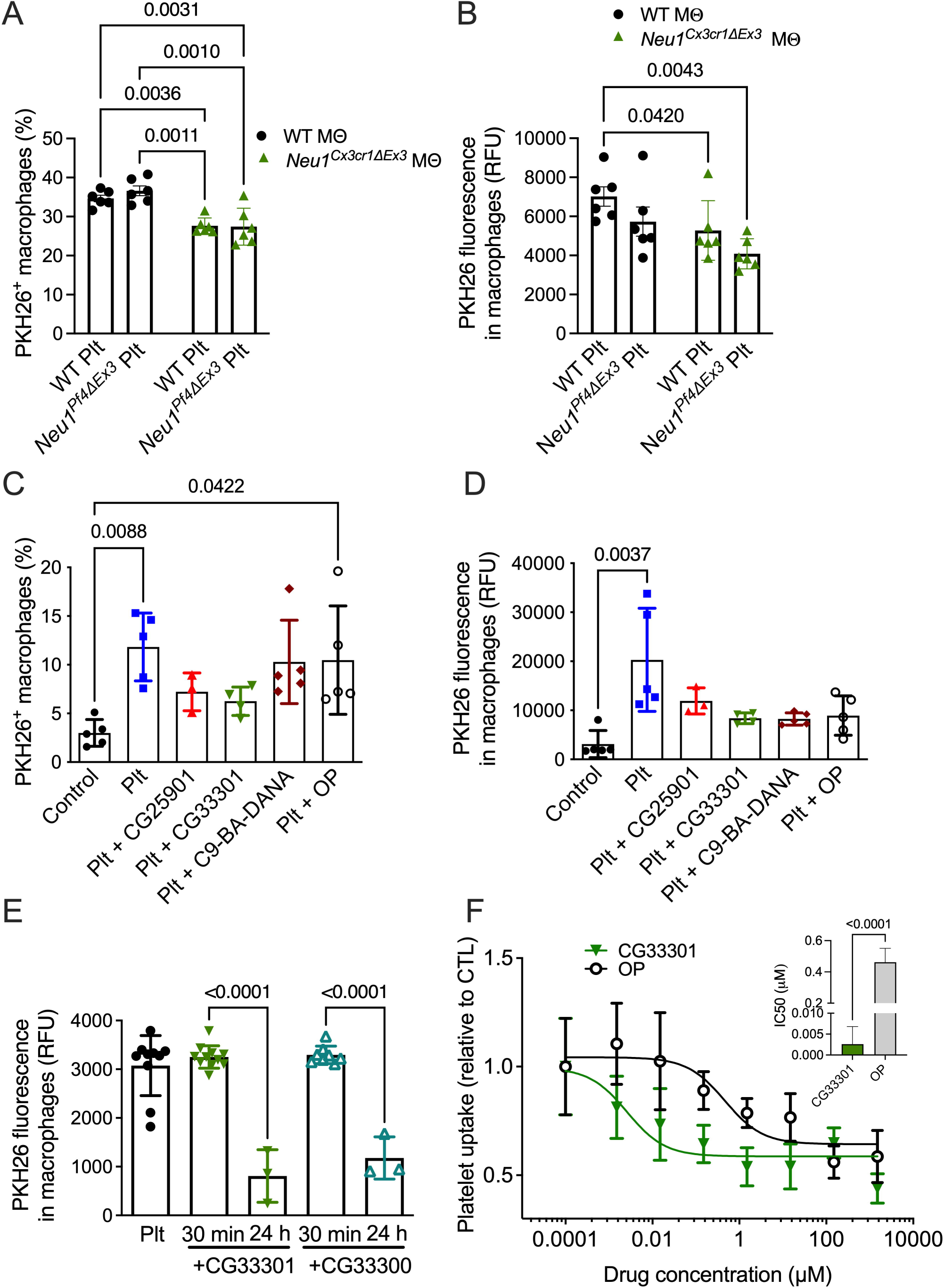
Macrophage NEU1 mediates platelet uptake in vitro. (A-B) An uptake of PKH26-labeled WT or platelet KO (*Neu1^Pf4^*^Δ*Ex3*^) anti-CD41a opsonized platelets by WT or macrophage KO (*Neu1^Cx3cr1^*^Δ*Ex3*^) cultured macrophages was analyzed by flow cytometry. **(A)** Fraction (%) of CD11b^+^ macrophages positive for PKH26 fluorescence. **(B)** Mean fluorescence intensity of PKH26^+^ macrophages, reflecting the number of platelets ingested per macrophage. Genetic deficiency of macrophage NEU1, but not platelet NEU1, significantly reduces platelet uptake. **(C-D)** Pretreatment of macrophages with the specific NEU1 inhibitors but not OP reduces platelet uptake measured as fraction **(C)** or mean fluorescent intensity **(D)** of PKH26^+^ macrophages. **(E)** Platelet uptake is inhibited by a 24 h pre-treatment of macrophages with specific NEU1 inhibitor CG33300 or its esterified form CG33301. Both compounds added to the macrophage cultures immediately before addition of platelets do not inhibit the uptake. **(F)** Concentration dependence of the platelet uptake inhibition by OP and CG33301. Insert shows a comparison of IC50 values for CG33301 and OP. Viability of cells measured with Zombie Aqua™ dye in all experiments exceeded 90%. Graphs show individual values, means and SD (A-E) or means and SD (F) of 3-8 biological replicates (individual cultures). *P* values were calculated by one-way ANOVA with Tukey post hoc test (**A-E**) or *t* test **(F)**. Only *P* values <0.05 are shown.

Hepatocytes and liver macrophages (Kupffer cells) were reported to play the major role in depletion of desialylated platelets via Ashwell-Morell receptor (ASGPR1/2) [7–9, 11]. To study the role of NEU1 in this process, we have analyzed the the uptake of CD41-opsonised platelets by primary hepatocytes and Kupffer cells isolated either from WT or NEU1-deficient mice. These experiments demonstrated that Kupffer cells from NEU1-deficient mice showed somewhat lower uptake rates of WT platelets as compared to Kupffer cells from the WT mice (Fig. S5), while the hepatocytes from both WT and NEU1-deficient mice showed a similar uptake rate (Fig. S6). In contrast, Kupffer cells showed a drastically reduced uptake of platelets isolated from NEU1 deficient mice compared to WT platelets, confirming that NEU1 is essential for platelet clearance by liver macrophages (Fig. S5).

We further tested whether pharmacological neuraminidase inhibitors could also inhibit the platelet uptake by spleen macrophages. We used previously described specific inhibitor of NEU1 C9-BA-DANA (IC_50_ 3,400 nM [42]) or highly specific NEU1 inhibitors CG33300 (IC_50_ 140 nM) and CG25900 (IC_50_ 420 nM) developed by our team (see ref. [43] and Supplemental methods). The latter two compounds were used as their C1-methyl ester forms (CG33301 and CG25901, respectively) with improved LogP values making them more suitable for in cellulo/in vivo experiments[43, 44]. Both compounds act as prodrugs, with conversion to the active acid forms by cellular esterases [45, 46]. We also tested an influenza virus neuraminidase inhibitor, oseltamivir phosphate (OP), that shows only a limited activity against NEU1 (IC_50_ ∼500 μM [22, 23]) but was previously reported to improve platelet counts in some ITP patients. In the first experiment, to block phagocytic capacity of macrophages, the inhibitors were added to the cultured macrophages 24 h prior to loading opsonized platelets at 15 μM concentrations, which exceeds 5xIC_50_ for each inhibitor except for OP. We found that a 24 h pretreatment of macrophages with C9-BA-DANA, CG33301 and CG25901, but not with OP, reduced the platelet uptake measured either as a fraction of PKH26^+^ macrophages or as the mean fluorescence intensity per cell (Fig. 3E-F). In contrast, no inhibition of the platelet uptake was detected when the specific NEU1 inhibitor CG33300 or its esterified form CG33301 were added to the culture medium immediately before platelets to prevent surface desialylation of platelets either by platelet or macrophage NEU1 during the 30-min uptake window (Fig. 3G). These results are consistent with the hypothesis that the NEU1 inhibitors blocked engulfment of opsonised platelets by splenic macrophages by interfering with the phagocytic activity of the latter cells but not by blocking desialylation of the platelet surface. We further analysed the concentration dependence of the platelet uptake inhibition by OP and the most potent NEU1 inhibitor, CG33301. Both compounds were added to the macrophage cultures 24 h before platelets in the range of concentrations between 1,500 μM and 0.00015 μM. Although OP could inhibit the platelet uptake at concentrations above 150 μM, CG33301 showed ∼200-fold increase in potency compared to OP (IC_50_ 0.004 μM for CG33300 vs 0.9 μM for OP, p<0.007) (Fig. 3G).

### Pharmacological inhibition of NEU1 rescues platelet depletion in the mouse model of ITP

We assessed whether the administration of the NEU1 inhibitor CG33301 could mitigate thrombocytopenia *in vivo* in the passive mouse ITP model. A preliminary pharmacokinetic analysis demonstrated that after IP injection in mice, inhibitors similar to CG25901 were rapidly converted to the active form, reaching maximal concentration in the blood approximately 0.5 h after the administration (*T_1/2_* in the blood was approximately 1.8 h for the active and 4 h for the methyl ester; Cairo, C.W., unpublished). Based on these results, we administered CG33301 twice, 1 h before and 2 h after the injection of the platelet-depleting anti-CD41a antibodies which we expected to result in a steady concentration of the inhibitor in the blood within the 6 h time window crucial for the platelet depletion (see Fig. S2 for experimental design). We, first, tested CG33301 in the dose of 30 mg/kg BW, previously shown to result in complete inhibition of the endogenous NEU1 in the mouse tissues [12] (Fig. 4A). In a separate experiment, we compared the action of CG33301 and OP using the lower dose of 2 mg/kg BW and the same regimen for both drugs (Fig. 4B). Treatment with the specific NEU1 inhibitor CG33301 preserved platelet counts at both 2 mg/kg BW and 30 mg/kg BW doses (Fig. 4A, B). At 6 h post anti-CD41a antibody injection, mice receiving CG33301 at 2 mg/kg BW maintained significantly higher (*p*<0.01) platelet counts compared to untreated mice or mice treated with OP (Fig. 4A,B). Moreover, the platelet counts in mice treated with CG33301 at 30 mg/kg BW were not statistically different from those in mice that did not receive the anti-CD41a antibody (Fig. 4A). Mice treated with OP in the dose of 2 mg/kg BW showed a drop in platelet counts similar to that of untreated mice (Fig.4B)

**Figure 4.**
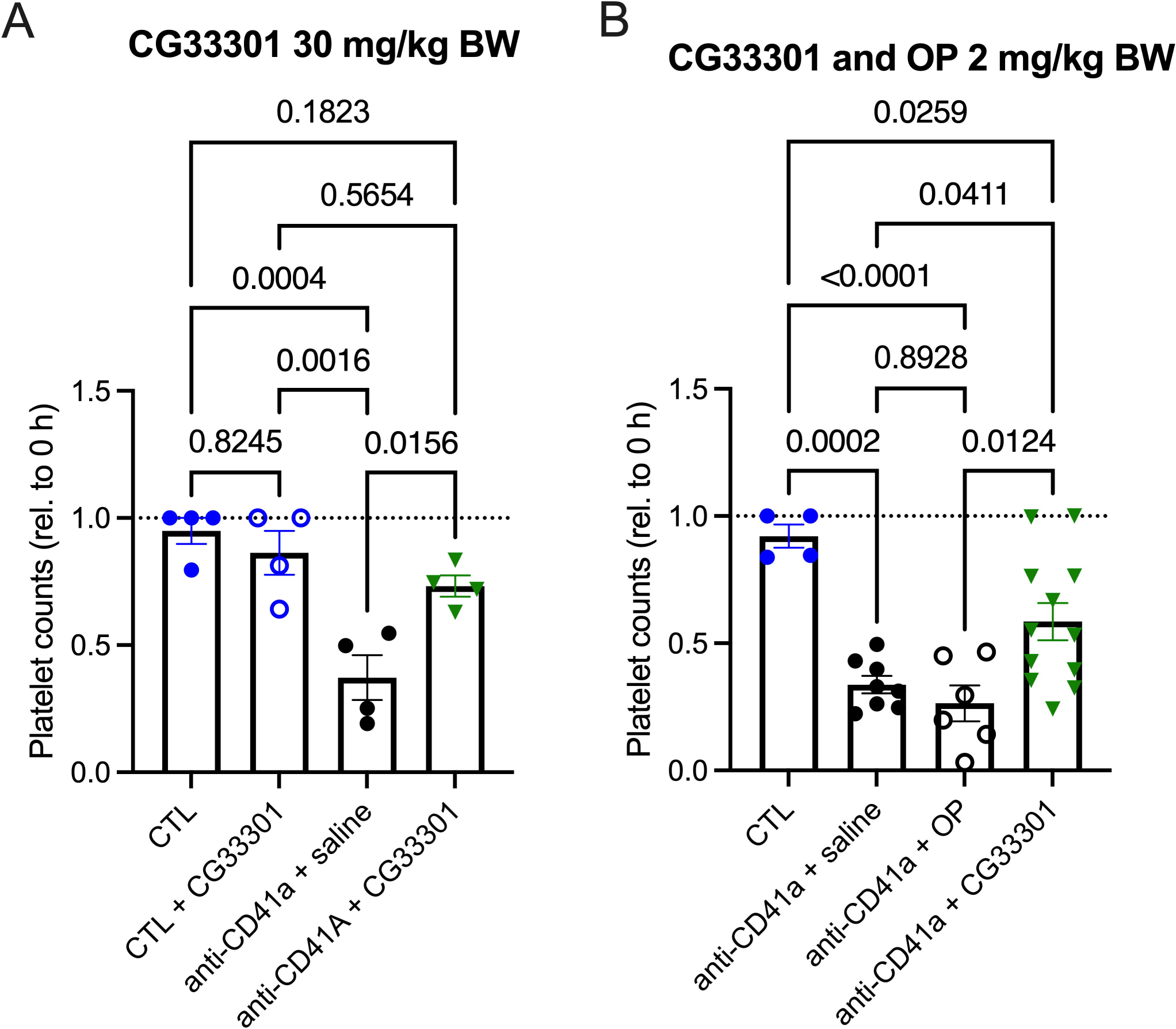
Pharmacological inhibition of NEU1 protects against ITP *in vivo*. WT animals received IP injections of neuraminidase inhibitors in saline or saline only 1 h before and 2 h after the injection of anti-CD41a antibodies to induce platelet clearance. Blood was sampled from the mandibular vein at 0 h and 6 h. Platelet counts were measured by FACS and shown as fold change from 0 h. **(A)** Platelet levels in the ITP mice, treated with CG33301 at the dose of 30 mg/kg BW, are increased compared to mice treated with vehicle (saline) and similar to untreated mice (CNT) or mice treated with CG33301 only (CNT + CG33301). **(B)** At the dose of 2 mg/kg, CG33301, but not OP preserves platelet counts in the ITP mouse model. Graphs show individual data and means ± SEM, n=4–11/group. *P* values were calculated using one-way ANOVA with Tukey post-hoc test.

## Discussion

In this study, we demonstrate that macrophage, but not platelet NEU1, is a critical driver of platelet clearance by splenocytes in immune thrombocytopenia. Using conditional NEU1 KO mouse strains, in vitro platelet uptake assays, and selective neuraminidase inhibitors, we show that the genetic inactivation or pharmacological blockage of NEU1 in spleen macrophages protects against thrombocytopenia in the passive mouse ITP model. These findings establish NEU1 as a novel therapeutic target in ITP and highlight the importance of macrophage sialoglycoconjugate homeostasis in platelet turnover.

While the classical paradigm of ITP emphasizes autoantibody-mediated platelet opsonization and Fc receptors for immunoglobulin G (Fcγ)–dependent phagocytosis by splenic macrophages, accumulating evidence has revealed additional Fc-independent platelet clearance pathways involving, in particular, desialylation of platelet surface proteins. Desialylated platelet proteins expose underlying galactose residues, rendering them susceptible to recognition by the asialoglycoprotein (Ashwell-Morell) receptor and facilitating their clearance by Kupffer cells and hepatocytes [9, 10]. Other lectin receptors, including MGL, CLEC4F, and CD11b, have been also implicated in the glycan-mediated platelet clearance [8, 47–49]. In turn, platelet desialylation was assumed to be conducted by NEU1 at the platelet surface in response to the stimuli induced by auto-antibodies, platelet activation or inflammation.

Multiple platelet glycoproteins are targeted by autoantibodies in ITP. The most frequent targets are GPIIb/IIIa, GPIb/IX and, less frequently, GPIa/IIa and GPVI [50]. The effect on platelets in terms of activation, desialylation, and apoptosis may depend on antibody specificity, but this remains unclear [51–53]. For example, anti GPIIb/IIIa antibodies more readily induce platelet desialylation, whereas anti GPIb/IX antibodies may drive more pronounced apoptotic signaling [19]. Platelets express the low-affinity IgG receptor FcγRIIA, which could be activated by some autoantibodies [54]. FcγRIIA engagement can trigger platelet activation, but also desialylation and apoptosis, contributing to accelerated platelet clearance and thrombocytopenia [55]. However, a consistent correlation between a given glycoprotein target and a particular pattern of platelet dysfunction has not been firmly established in clinical practice [56]. Our current results extend this model by showing that the macrophage NEU1 actively contributes to desialylation of platelets and activates platelet phagocytosis, while the platelet pool of NEU1 is not involved in the regulation of their clearance.

Our study demonstrates a drastic difference in the basal platelet levels and the response to administration of anti-platelet antibodies for the constitutive and tissue-specific NEU1 knockout mouse models. Macrophage-specific or platelet-specific NEU1 deletion had no effect on basal platelet levels. The reduced basal platelet counts observed in constitutive *Neu1* KO mice likely reflect systemic consequences of lifelong NEU1 deficiency rather than a direct platelet-intrinsic effect. NEU1-null animals develop progressive inflammatory and metabolic abnormalities, splenomegaly, and altered tissue homeostasis, all of which may affect megakaryopoiesis and platelet survival. In contrast, the absence of thrombocytopenia in macrophage– or platelet-specific NEU1 depletion indicates that the enzyme does not regulate steady-state platelet homeostasis in a cell-autonomous manner but rather modulates pathological platelet clearance during immune-mediated conditions [25, 30, 31, 57–60]. Notably, alterations in the thrombopoiesis have been previously reported in the mouse strains with depleted glycosyltransferases and altered glycosylation patterns [61].

Platelet-specific NEU1 KO mice showed levels of thrombocytopenia similar to those of WT mice after ITP induction by either anti-CD41a or anti-CD42b antibodies, the latter are reported to trigger the removal of sialic acids from GPIbα protein by activating NEU1. Their platelets also showed surface sialylation levels similar to those of WT mice. By contrast, in macrophage-specific and constitutive NEU1 KO mice as well and in the galactosialidosis mice with 90% of NEU1 deficiency the platelet deletion rate was significantly reduced and platelets from NEU1-deficient mice showed an excessive sialylation. These results were supported by in vitro experiments showing that the macrophages from NEU1-deficient mice, or those pretreated with a specific NEU1 inhibitor, showed a lower opsonized platelet uptake compared to WT cells. In contrast, the uptake rates for WT and NEU1-deficient platelets were similar. Notably, the uptake rate of platelets for NEU1-deficient platelets by liver Kupffer cells and hepatocytes was drastically reduced compared to the platelets from WT mice. Together, these results indicate that while the platelet NEU1 is essential for their desialylation-dependent clearance by liver, in the spleen, macrophage, rather than platelet NEU1, orchestrates the depletion of antibody-opsonised thrombocytes.

Our results are also consistent with earlier work from our group as well as from other laboratories showing that NEU1 translocates from the lysosomes to the plasma membrane during the differentiation of macrophages, where it desialylates receptors that trigger macrophage activation, cytokine secretion and phagocytosis[25, 28, 30, 62–70]. In particular, NEU1-deficient macrophages and immature dendritic cells showed impaired capacity to engulf IgG-opsonized beads and erythrocytes, while their binding measured at 4 °C was not affected [30]. Notably, FcγR receptors on the surface of NEU1-deficient macrophages responsible for the uptake of opsonised cells were hypersialylated. Our results also revealed an impaired FcγR phosphorylation as well as markedly reduced phosphorylation of the downstream Syk kinase in response to treatment of macrophages with IgG-opsonized beads suggesting that NEU1 activates the phagocytosis in macrophages by desialylation of surface receptors. Our current data provide evidence that the macrophage NEU1 is also critical for platelet clearance in ITP.

Currently available treatments for ITP include corticosteroids and intravenous immunoglobulin, as well as more targeted therapies including rituximab, and thrombopoietin receptor agonists [71, 72]. Still, many patients relapse or require long-term or multimodal therapies with associated side effects. Repurposing viral neuraminidase inhibitors such as OP, approved for treatment of influenza, has been proposed as a novel strategy and tested in murine ITP models and in open-labeled trial with limited number of patients. Although OP has been shown to improve platelet counts in some refractory patients [19], the effect was limited, which is not surprising considering that the drug primarily targets viral neuraminidases and is not specific for NEU1. In our experiments, the selective NEU1 inhibitor CG33301 outperformed OP in preserving platelet counts during ITP induction in mice. The improved potency of CG33301 over OP is likely due to the specificity of the former compound for the target, NEU1. In contrast, given the weak inhibitory activity of OP toward mammalian NEU1, any clinical benefit of this agent in ITP is likely indirect or mediated through off-target effects rather than through the specific NEU1 blockage. This, in our opinion, supports further research aimed on the development of bioavailable highly selective NEU1 inhibitors as therapeutic candidates for ITP. Importantly, while acute NEU1 inhibition is protective, chronic pan-tissue depletion of NEU1 (such as in constitutive *Neu1* knockout mice) reduces baseline platelet counts, suggesting that transient pharmacologic inhibition of the enzyme may be preferable to long-term suppression.

Overall, our findings resonate with the emerging concept that the protein and cellular glycosylation/sialylation axis is central to hematologic immune regulation. Sialic acids and neuraminidases modulate not only platelet clearance but also receptor signaling, immune cell activation, and inflammatory responses. Therapeutically, targeting NEU1 may have broader effects beyond ITP, including potential applications in autoimmune cytopenias, sepsis-associated thrombocytopenia, or transfusion medicine where storage-related desialylation compromises platelet viability. Future studies should extend these findings to human macrophages from ITP patients, assessing whether NEU1 activity correlates with disease severity or therapeutic response. In addition, clinical translation of selective NEU1 inhibitors will require careful monitoring of their potential off-target effects, given NEU1’s pleiotropic role in lysosomal catabolism, connective tissue homeostasis, and receptor signaling (reviewed in [29, 73–76]). Nevertheless, our results provide a rationale for future use of selective NEU1 inhibitors as novel therapeutics for ITP and related thrombocytopenic disorders.

## Acknowledgements

The authors thank Dr. Mila Ashmarina for critical reading of the manuscript.

## Availability of data and materials

The data, analytic methods, and study materials will be made available to other researchers for purposes of reproducing the results or replicating the procedure.

## Author’s contribution

Conducted experiments: AC, DP, SN, LC, XP, MS; analysed data: AC, LC, DP, SN, TP; conceptualization and funding: AVP, CC, YP, RB, TP; manuscript preparation: (first draft) AC and AVP; manuscript preparation (editing): TP, YP, RB, CC, AVP.

