## Supplemental figures and tables for "Inhibition of macrophage neuraminidase 1 protects against immune thrombocytopenia by limiting platelet clearance"

**Table S1: Antibodies used in the study and their working concentrations**

| Marker | Fluorochrome | Clone (manufacturer) | Amount / test | Final concentration |
|---|---|---|---|---|
| CD45 | BV785 | 30-F11 (BD) | 0.25 µg / test | ~2.5 µg/mL |
| CD11b | PerCP-Cy5.5 | M1/70 (BD) | 0.25–0.5 µg / test | 2.5–5 µg/mL |
| CX3CR1 | BV605 | SA011F11 (Biolegend) | 0.25 µg / test | ~2.5 µg/mL |
| CD41 | APC | MWReg30 (BD) | 0.25 µg / test | ~2.5 µg/mL |
| Ter119 (Ter-119) | FITC | TER-119 (BD) | 0.25 µg / test | ~2.5 µg/mL |
| Fc block (CD16/32) | — | 2.4G2 (BD) | 0.5–1 µg / test | 5–10 µg/mL |
| F4/80 | APC/Fire 750 | BM8 (BioLegend) | 0.25 µg / test | ~2.5 µg/mL |
| ASGPR1 | AF647 | 8D7 (Santa Cruz Bio) | 0.25 µg / test | ~2.5 µg/mL |
| SNA lectin | Cy5 | EBL (Vector Lab) | 0.25 µg / test | ~2.5 µg/mL |
| WGA lectin | FITC | EBL (Vector Lab) | 0.25 µg / test | ~2.5 µg/mL |
| PNA lectin | Cy5 | EBL (Vector Lab) | 0.25 µg / test | ~2.5 µg/mL |
| CD42d* | APC | 1C2 (BioLegend) | 0.25 µg / test | ~2.5 µg/mL |
| CD62P* | FITC | AK4 (BioLegend) | 0.25 µg / test | ~2.5 µg/mL |

*Antibodies were used for internal control of platelets purity and activation.

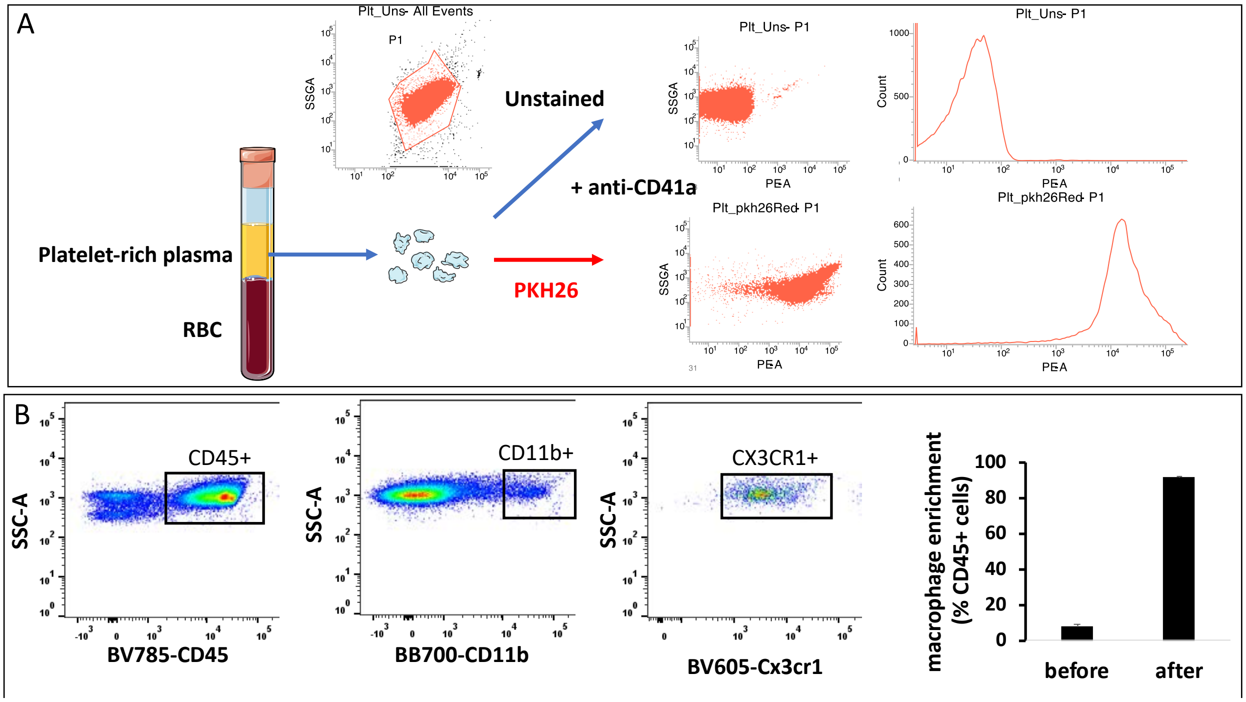


**Supplementary Figure S1. Experimental design and gating strategy for in vitro platelet uptake assay by spleen macrophages.**

**A.** Representative flow cytometry plots showing discrimination of unlabelled and PKH26-labeled platelets. The cell viability, assessed by Zombie Aqua™ dye (BioLegend) always acceded 92%.
**B.** Gating strategy for splenic macrophages. $CD45^{+}CD11b^{+}CX3CR1^{+}$ isolated population was used for platelet uptake analysis. Graph shows a percentage of macrophages in a total population of CD45+ cells before and after macrophage isolation. N=23

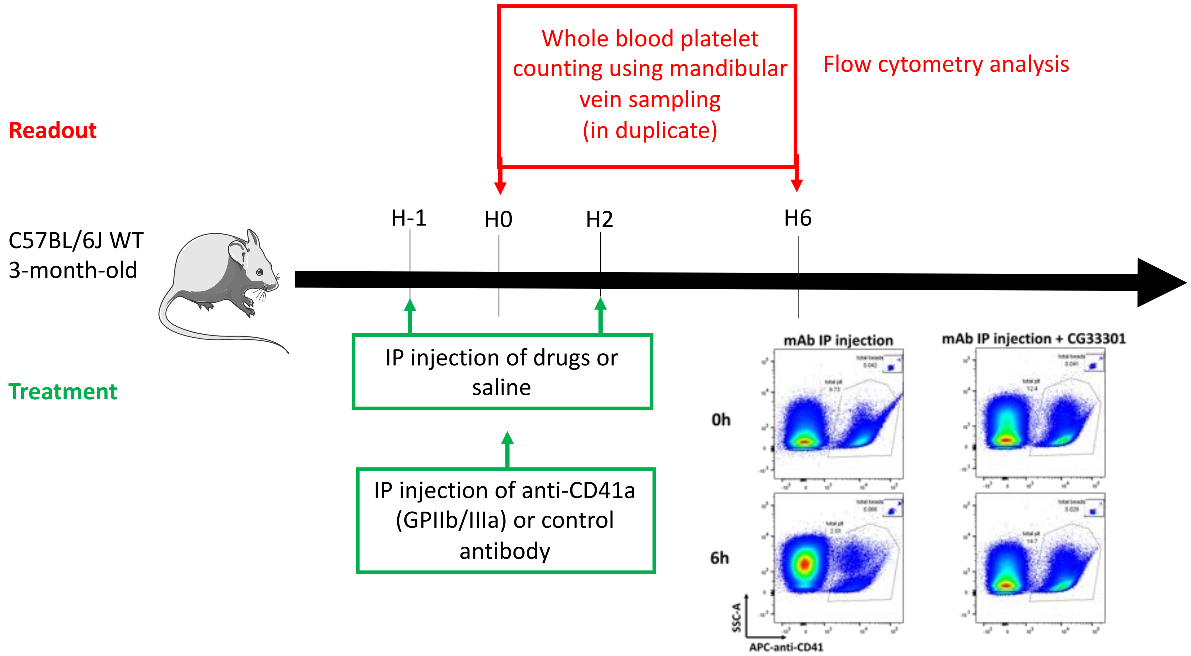


**Supplementary Figure S2. Experimental design for testing the efficacy of pharmacological neuraminidase inhibitors in passive mouse ITP model.**

Three-month-old C57BL/6J mice received intraperitoneal injection of anti-CD41a (GPIIb/IIIa) antibody or control mouse IgG. Neuraminidase inhibitors (CG33301 or oseltamivir phosphate) in saline or saline only were administered intraperitoneally one hour before and two hours after administration of the antibody. After 6 hours, platelets were counted by flow cytometry in the whole blood collected from the mandibular vein.

Insert shows a representative flow cytometry results demonstrating that CG33301 in the dose of 30 mg/kg BW protects against ITP-induced platelet depletion.

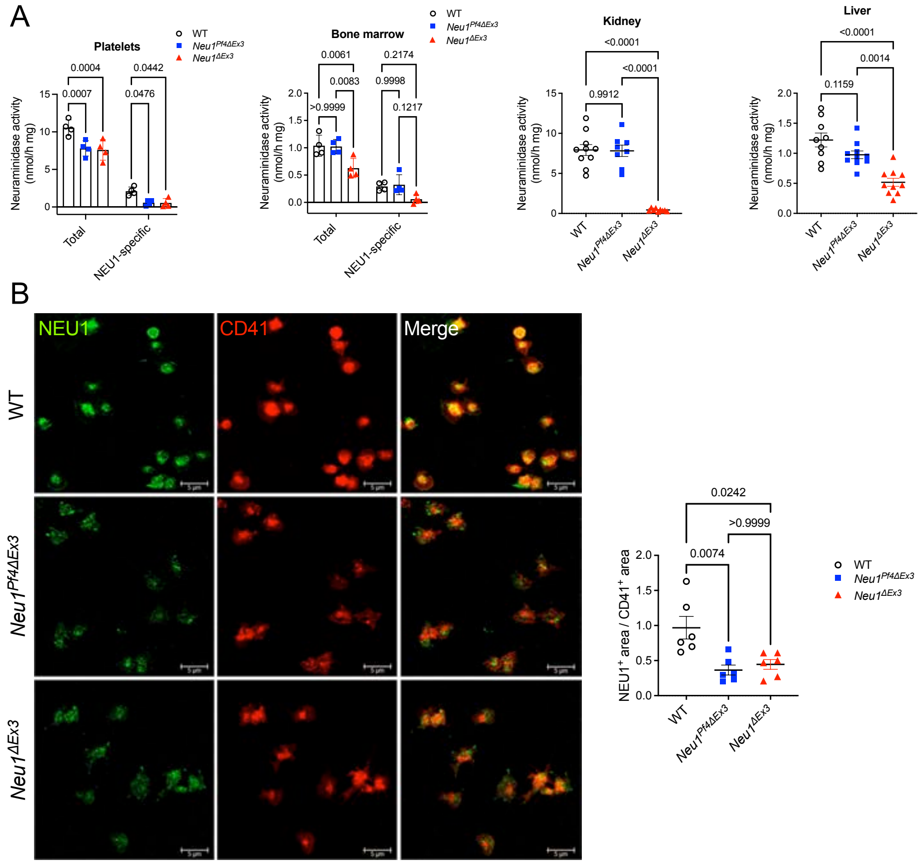


**Supplementary Figure S3. Platelets of *Neu1$^{Pf4\Delta Ex3}$* mice show reduced total and NEU1-specific neuraminidase activity and reduced labeling with anti-NEU1 antibodies.**

**A.** Total acidic neuraminidase (NEU1, NEU3 and NEU4 together) activity was measured in the purified platelets, bone marrow, kidney and liver homogenates from WT, *Neu1$^{Pf4\Delta Ex3}$* and constitutive KO *Neu1$^{\Delta Ex3}$* mice using the fluorogenic 4-MU-NANA substrate at pH 4.6. To measure NEU1-specific activity, the platelet and bone marrow homogenates were supplemented with the specific NEU3/NEU4 inhibitor GLT 1-77. NEU2-specific activity in the platelet homogenates was measured at pH 7.5. Both total and NEU1-specific neuraminidase activities were reduced in platelets of *Neu1$^{Pf4\Delta Ex3}$* compared to WT mice. In contrast the neuraminidase activity in the bone marrow, kidney and liver of *Neu1$^{Pf4\Delta Ex3}$* was similar to that WT mice. *Neu1$^{\Delta Ex3}$* mice showed reduced neuraminidase activity in all studied tissues. Individual results, mean and SEM are shown. *N* = 4 (platelets and bone marrow) or *N* = 8-10 (liver and kidney); *P* values were calculated by one-way ANOVA with Tukey post-hoc test.

**B.** Purified platelets from WT, *Neu1$^{Pf4\Delta Ex3}$* *Neu1$^{\Delta Ex3}$* mice were fixed, permeabilized and labeled with antibodies against mouse NEU1 (green) and platelet marker CD41 (red). Panels show representative images captured using Leica DM5500 Q upright confocal microscope (63x oil objective, N.A. 1.4; 5x digital Zoom). Size bars equal 5 µm. Bar graph shows quantification of fluorescence areas with ImageJ software. Individual results, mean and SEM are shown. *N* = 6; *P* values were calculated by one-way ANOVA with Tukey post-hoc test.

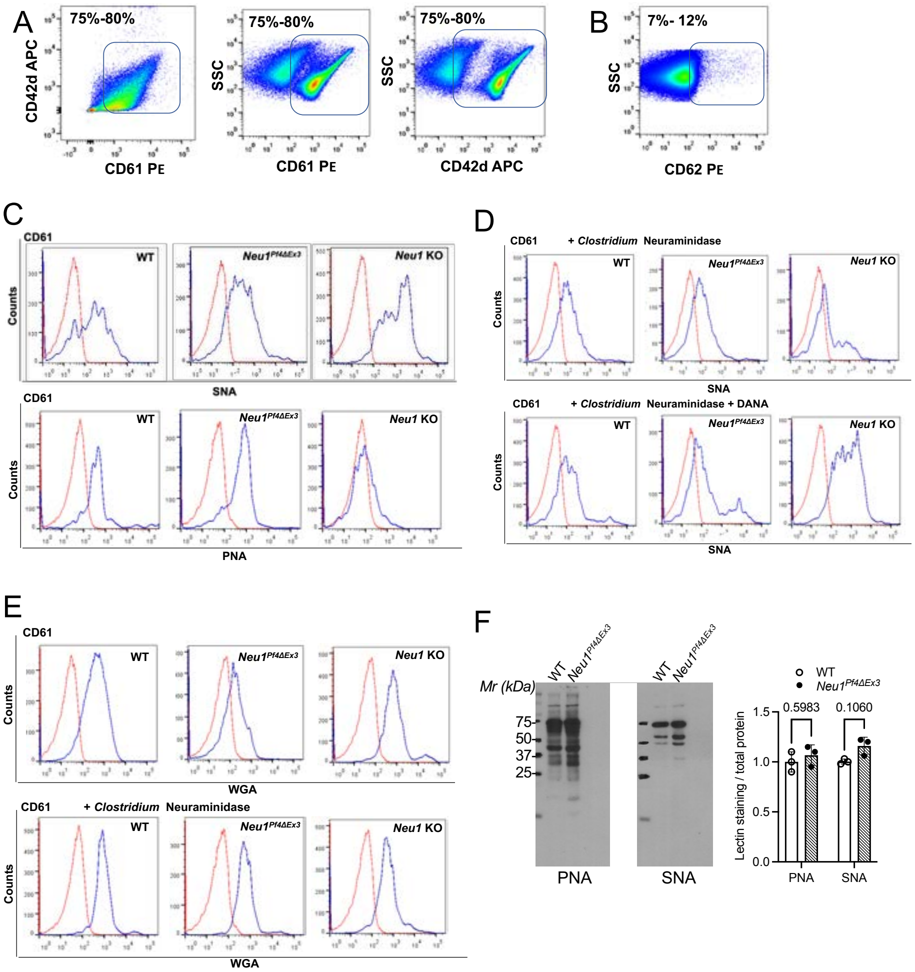


**Supplementary Figure S4. Platelets of *Neu1$^{Pf4\Delta Ex3}$* mice show sialylation profiles similar to those of WT mice.**

**(A-B)** Gating strategy for purified mouse platelets. Representative flow cytometry plots showing that 75-80% of analysed cells were positive for the platelet markers CD61 and CD42 (A). Only 7-12% of cells expressed the marker of activated platelets CD62. **(C-E)** Flow cytometry histograms of the platelets purified from the blood of WT, *Neu1$^{Pf4\Delta Ex3}$* and constitutive *Neu1* labeled with fluorescent anti-CD61 and antibodies (red traces) and lectins (blue traces) specific for sialic acids (SNA), desialylated terminal galactose residues (PNA) and N-acetyl-D-glucosamine (WGA) as indicated. **(C)** Platelets from WT and *Neu1$^{Pf4\Delta Ex3}$* mice show similar affinity to SNA and PNA, while platelets from *Neu1* KO mice show increased SNA and reduced PNA labeling consistent with their increased sialylation. **(D)** Treatment with an assess of bacterial neuraminidase reduces their labeling by SNA while treatment with neuraminidase in the presence of pan-neuraminidase inhibitor DANA does not change SNA levels. **(E)** WGA labeling is similar for platelets from WT and *Neu1$^{Pf4\Delta Ex3}$* and *Neu1* KO mice and is not changed by neuraminidase treatment. All panels show representative histograms of the analyses performed in triplicate with cells purified from the pooled blood of 5 mice per genotype. **(F)** Representative lectin blots of platelets from WT and *Neu1$^{Pf4\Delta Ex3}$* mice labeled with biotinylated PNA and SNA lectins and quantification of total band intensity per lane with ImageJ software. Graphs show individual results, means and SD for 3 mice/genotype. P values were calculated by two-way ANOVA with Sidak post hoc test.

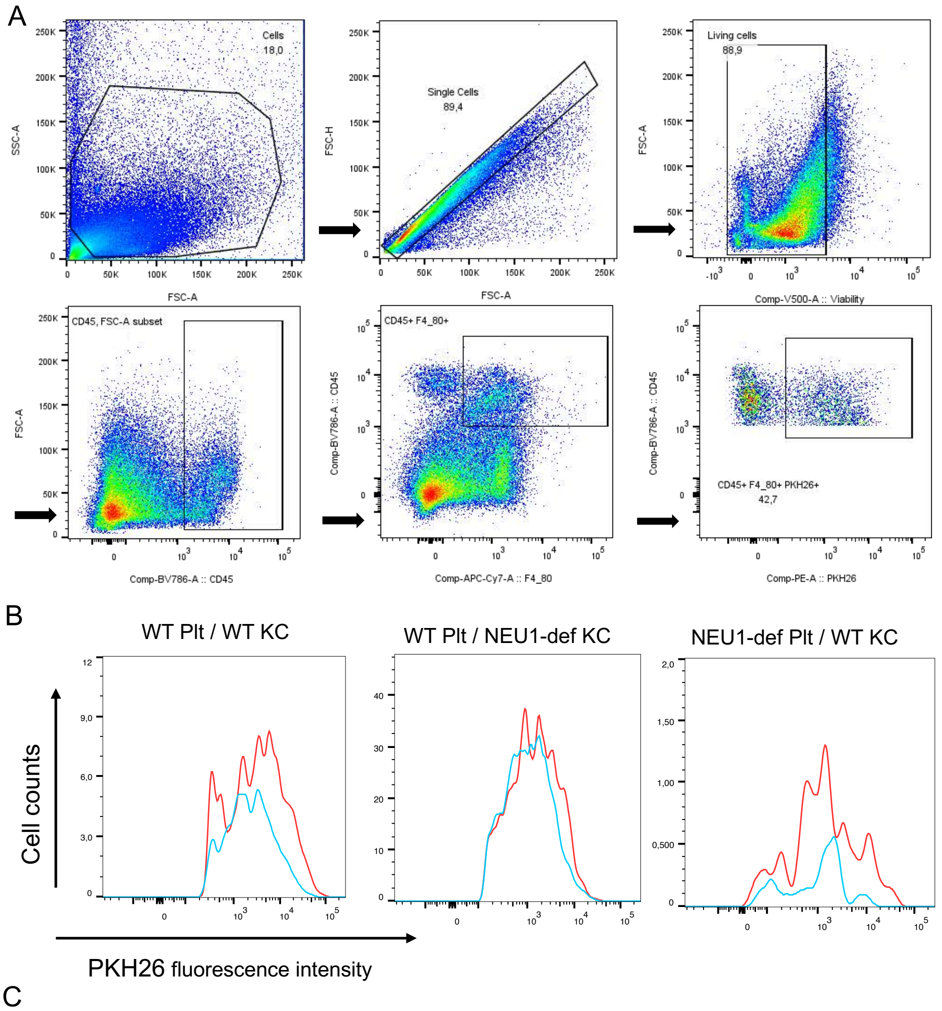


C

| | WT Plt / WT KC | | WT Plt / NEU1-def KC | | NEU1-def Plt / WT KC | |
|---|---|---|---|---|---|---|
| | **4ºC** | **37ºC** | **4ºC** | **37ºC** | **4ºC** | **37ºC** |
| **CD45⁺ F4/80⁺ PKH26⁺ cells (%)** | 16 | 34.9 | 28.2 | 42.7 | 1.5 | 6.61 |
| **Mean PKH26 fluor. intensity** | 4287 | 7010 | 2071 | 2358 | 1874 | 3900 |

**Supplementary Figure S5. Uptake of platelets by cultured primary mouse Kupffer cells.** Primary Kupffer cells (KC) isolated from WT and NEU1-deficient mice were incubated for 30 min at 37 °C or 4 °C with PKH26-labeled WT or Neu1-deficient platelets (Plt). **(A)** Gating strategy used to identify viable Kupffer cells (CD45$^+$F4/80$^+$). **(B)** Representative histograms showing PKH26 fluorescence intensity (x-axis) versus cell count (y-axis) in gated CD45$^+$F4/80$^+$ KC population following incubation with PKH26-labeled anti-CD41a IgG-opsonises platelets at 37 °C (red traces) or 4 °C (blue traces). **(C)** Percentage of PKH26$^+$ KC within the CD45$^+$F4/80$^+$ population and the mean PKH26 fluorescence intensity for each experimental condition. The cell viability, assessed by Zombie Aqua™ dye (BioLegend) always acceded 88%.

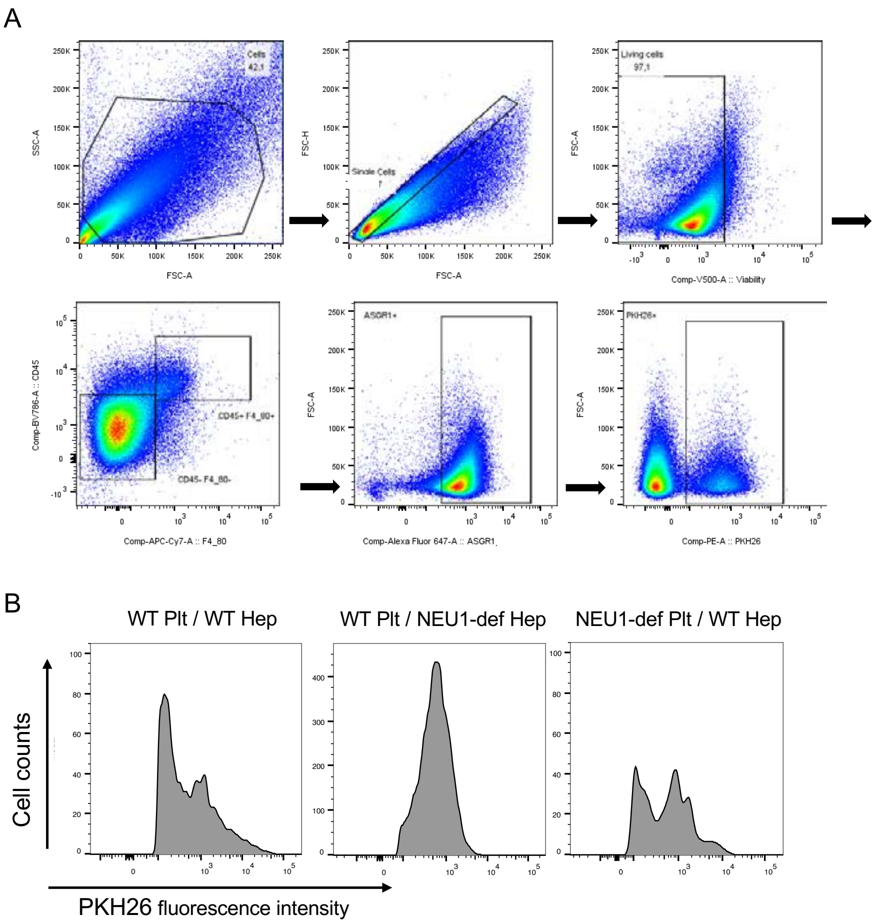


C

| | WT Plt / WT Hep | WT Plt / NEU1-def Hep | NEU1-def Plt / WT Hep |
|---|---|---|---|
| **$ASGPR1^{+}$/$PKH26^{+}$ cells (%)** | 13.8 | 11.8 | 9.32 |
| **PKH26 MFI** | 1826 | 759 | 1160 |

**Supplementary Figure S6. Uptake of platelets by primary cultured mouse hepatocytes.** Primary hepatocytes (Hep) isolated from WT and Neu1-deficient (Neu1-def) mice were incubated for 30 min at 37 °C with PKH26-labeled WT or Neu1-deficient platelets (Plt). **(A)** Gating strategy used to identify hepatocytes ($CD45^{-}$/$F4/80^{-}$/$ASGPR1^{+}$). **(B)** Representative histograms showing PKH26 fluorescence intensity (x-axis) versus cell counts (y-axis) in gated $CD45^{-}$/$F4/80^{-}$/$ASGPR1^{+}$ hepatocytes following incubation with PKH26-labeled anti-CD41a IgG-opsonises platelets at 37 °C. **(C)** Percentage of $PKH26^{+}$ hepatocytes within the $ASGPR1^{+}$ population and the mean PKH26 fluorescence intensity for each experimental condition. The cell viability, assessed by Zombie Aqua™ dye (BioLegend) always acceded 95%.

### Supplemental Materials and Methods.

#### Animals

The constitutive *Neu1* KO (*Neu1$^{\Delta Ex3}$*), the phagocyte-specific conditional *Neu1* KO (*Neu1$^{Cx3cr1\Delta Ex3}$*) and the cathepsin A hypomorph galactosialidosis (*CathA$^{S109A\text{-}Neo}$*) mouse models were previously described [1, 2]. A platelet-specific conditional *Neu1* KO (*Neu1$^{Pf4\Delta Ex3}$*) mouse was generated by crossing previously described *Neu1$^{loxPEx3}$* strain with the *Neu1* exon 3, flanked with the loxP sites [1], with the C57BL/6-$^{Tg(Pf4\text{-}icre)Q3Rsko}$/J strain (The Jackson Laboratory strain 008535), expressing the Cre recombinase under the control of the *Pf4* (platelet factor 4) gene promoter. *Neu1$^{Pf4\Delta Ex33}$* homozygous mice were compared with age- and sex-matched *Neu1$^{loxPEx3}$* mice. To generate inducible *Neu1 KO* mice, *Neu1$^{loxPEx3}$* strain was crossed with B6.Cg-*Ndor1$^{Tg(UBC\text{-}cre/ERT2)1Ejb}$*/1J strain (The Jackson Laboratory strain 007001) expressing the Cre-ERT2 fusion gene under the control of the human ubiquitin C (UBC) promoter. Mice, homozygous for the *Neu1$^{loxPEx3}$* allele and homozygous or heterozygous for the *Ndor1$^{Tg(UBC\text{-}cre/ERT2)1Ejb}$* allele (*Neu1$^{\Delta Ex3\text{-}ind}$* mice), do not show NEU1 deficiency and are viable and fertile. Upon the treatment with tamoxifen (5 IP injections at 75 mg/kg BW over 5 consecutive days), they express Cre activity resulting in Cre-mediated recombination and deletion of the *loxP*-flanked exon 3 of the Neu1 gene in all studied cell and tissue types. All mice were on C56/BL6J inbreed background. Heterozygote breeding pairs were used to maintain the colony and generate the initial homozygous mutants, followed by homozygote breading to generate homozygous mice. Mice were housed in the Animal facilities of CHU Ste-Justine, following the guidelines of the Canadian Council on Animal Care (CCAC). The animals were kept in an enriched environment with a 12:12 h light/dark cycle, fixed temperature, humidity and continuous access to water and a normal chow diet (5% fat, 57% carbohydrate). Equal cohorts of male and female mice were studied separately for each experiment, and statistical methods were used to test whether the progression of the disease, levels of biomarkers or response to therapy were different for male and female animals. Since differences between sexes were not detected, the data for male and female mice were pooled together.

**Spleen macrophage isolation**

Macrophages were isolated from spleen of WT and *Neu1*$^{Cx3cr1\Delta Ex3}$ mice by forcing pieces of organs through a 100 µm nylon mesh cell strainer (BD Biosciences, Durham, NC, USA) pre-wet with RPMI 1640 media with the back of a 1 mL syringe plunger at 4°C. The cell strainer was washed with RPMI 1640 at 4°C to flush the cells through the nylon mesh. The two previous steps were repeated until only connective tissue remained in the cell strainer. Cells were centrifuged at 300 x *g* for 5 min at 4°C. Cells were resuspended in fresh RPMI 1640 and plated in 20 cm petri dishes. After 45 min, the non-adherent cells were removed, and the purity of macrophages was assessed by flow cytometry. Only the batches containing >85% of CD11b$^{+}$ cells with viability >90% were used in the experiments (Fig. S1B).

**Isolation and culture of mouse hepatocytes and Kupffer cells**

Mouse liver hepatocytes and Kupffer cells were isolated as described [3-6] with the following modifications. Mice were anesthetized and positioned on a dissection tray. Using sterile instruments, a midline laparotomy was performed, and the intestines were shifted to the right to expose the portal vein and vena cava. The liver was perfused by inserting a needle into the vena cava above the kidney with pre-warmed perfusion buffer (1× HBSS supplemented with 0.1% EDTA and 2.5% HEPES) until the liver was cleared of blood. Subsequently, 5 ml of pre-warmed digestion buffer (1× HBSS with 2.5% HEPES, 1 mg/mL Collagenase D, and 50 µg/mL DNase I) was transferred to the liver. After perfusion, the liver was dissected and the gallbladder was removed.

Approximately 200 mg of liver tissue was placed into individual Eppendorf tubes containing 1 mL of RPMI medium supplemented with 1 mg/mL of Collagenase D and 50 µg/mL of DNase I. The tissue was finely minced using scissors to facilitate digestion and incubated at 37°C for 30 minutes with shaking every 5–10 min to improve enzymatic digestion. The resulting digested cell suspension was filtered through a 70 µm cell strainer and rinsed with 20 mL of RPMI medium to optimize cell recovery. The cell suspension was centrifuged at 50 × *g* for 3 min at 4°C with low acceleration and low brake settings. Following centrifugation, the cell fractions were separated: the pellet contained the hepatocytes, while the supernatant contained the Kupffer cells.

For hepatocyte purification, 20 mL of DMEM plating medium was added to the pellet, and the cells were resuspended by gentle swirling. The suspension was centrifuged at 50 × *g* for 3 min at 4°C. Most of the supernatants were aspirated, leaving approximately 1 mL of fluid, and the cells were resuspended in 20 mL of DMEM plating medium before undergoing another centrifugation step at 50 × *g* for 3 min at 4°C. The supernatant was aspirated, and the cell pellet was resuspended in the appropriate culture medium (Williams' E medium supplemented with 1% glutamine and 1% penicillin-streptomycin solution). Following cell counting, the hepatocytes were seeded at a density of $10^5$ cells per well onto pre-prepared collagen-coated culture 6 well plate. The medium was gently changed after 3 to 4 h of incubation, and the experiments with the platelet uptake were initiated no later than the day following plating.

For Kupffer cell purification, the remaining supernatant was transferred to a new 50 mL conical tube and centrifuged at 600 × *g* for 10 min at 4°C. The supernatant was removed, and the cell pellet was resuspended in 20 mL of a 45% Percoll solution (9 mL of 100% Percoll mixed with

2 mL of 10× HBSS and 9 mL of $H_2O$). Next, 8 mL of 1× HBSS was carefully overlaid along the side of the tube. The gradient was centrifuged at 1400 × *g* for 30 mins at 4°C with low acceleration and low brake settings. The upper 10 mL of the gradient was collected into a new 50 mL tube, ensuring recovery of the entire cell band at the interface. Then, 20 mL of RPMI medium was added, and the suspension was centrifuged at 600 × *g* for 10 min at 4°C. The supernatant was discarded, and the cell pellet was resuspended in Gibco AIM V Medium. Following cell counting, the cells were seeded into 6-well plates at a density of $10^5$ cells per well. The culture medium was replaced after 30 min, after which the purified cells used in platelet uptake experiments.

**Platelet isolation**

Platelets were isolated from the whole blood of WT CD57Bl6J or $Neu1^{Pf4\Delta Ex3}$ mice as previously described [7]. Briefly, the blood, collected by terminal bleeding from the heart in sodium citrate microtainer (BD), was centrifuged at 200 x *g* for 8 min (no brake) at the room temperature (RT). After centrifugation, the upper layer without the buffy coat (platelet-rich plasma; PRP) was carefully aspirated and transferred to a 15 ml conical plastic tube containing 1 mL of Modified Tyrode's buffer (134 mM NaCl, 2.9 mM KCl, 0.34 mM $Na_2HPO_4$, 12 mM $NaHCO_3$, 20 mM HEPES, 1 mM $MgCl_2$, 5 mM Glucose, pH 7.3). After the centrifugation at 1000 x*g* for 5 min at RT, the supernatant was gently discarded, and the pellet was resuspended in modified Tyrode's buffer for subsequent experiments.

**Neuraminidase activity assays**

Isolated mouse platelets or dissected tissues were homogenized in water at a 1:3 (w/v) ratio (e.g., 100 mg of tissue in 300 μL of water) in Eppendorf tubes using a CLS-5001 cordless motor driven pestle. To measure total acidic neuraminidase activity, a reaction mixture was prepared containing 5 µL of tissue or cells homogenate, 12.5 µL of 0.1 M sodium acetate buffer, pH 4.6, 12.5 µL of 0.8 mM fluorogenic substrate 4-methylumbelliferyl-N-acetylneuraminic acid (Sigma-Aldrich) in $ddH_2O$ and 10 µL of $ddH_2O$. To measure NEU1 activity 10 µL of 0.5 mM NEU3/NEU4 specific inhibitor C9-4BPT-DANA (CG17700) [8] was added instead of 10 µL of $ddH_2O$. Neutral neuraminidase (NEU2) activity was measured similarly to the activity of total acidic neuraminidase but 12.5 µL of 0.1 M Tris buffer, pH 7.5 was added instead of the acetate buffer. After incubation at 37 °C for 1 h, the reaction was stopped by adding 210 µL of 0.4 M glycine buffer, pH 10.4. Then, 250 µL of the final reaction mixture was transferred to a black 96-well plate (CoStar), and the fluorescence of liberated 4-methylumbelliferone was measured using a ClarioStar plate reader (HSPCG Labtech). Protein concentration was determined using the Pierce BCA Protein Assay Kit (#23225, Thermo Scientific).

**Immunofluorescence microscopy**

Purified platelets sample ($10^6$ cells/slide) were resuspended in 100 µl of Tyrode buffer and loaded on Lys-coated microscope glass slides using Cytospin. Cells were allowed to dry before fixation with 4% paraformaldehyde at room temperature for 20 min. Cells were further washed three times with PBS, permeabilizes with 0.1% Triton X-100 in PBS for 10 min at room temperature and blocked with 5% BSA in PBS with for 2 h at room temperature. The cells were further labeled with primary antibodies against NEU1 (Abcam 244119; 1:100) and CD41a (Invitrogen 14-0411-82; 1:50) diluted in 1% BSA in PBS overnight at 4°C. Cells were further labeled with secondary antibodies Alexa Fluor 488 anti-rabbit IgG (1:200) and Alexa Fluor 555 anti-rat IgG (1:300) for 1 h at room temperature in the dark. Slides were further mounted with ProLong™ Gold Antifade medium (Invitrogen, P36935). Images were captured using a Leica DM5500 Q upright confocal microscope (63x oil objective, N.A. 1.4). Final figure panels were prepared using Adobe Photoshop.

**Flow cytometry**

Mouse and human cultured cells and mouse primary cells were labeled with a fixable viability dye eFluor® 506 (eBioscience, San Diego, CA, USA) in PBS. The cells of mouse origin were additionally treated with rat anti-mouse CD16/CD32 Fc receptor block (clone 2.4G2, BD Biosciences). The cells were further incubated with specific antibodies and fluorescently labeled lectins (see the Table S1 for antibodies and their working concentrations) in PBS supplemented with 5% of fetal bovine serum (FBS, qualified, Canada origin, Life Technologies, Burlington, Canada), fixed with 4% PFA in PBS during 15 min at 4°C, washed and analyzed by flow cytometry using a BD LSRFortessa cell analyzer (BD Biosciences). Fluorescence minus one (FMO) and/or isotype staining controls were used to adjust the gates and to determine the fluorescence background. Data analysis was performed using the FlowJo software (version 10.1, Tree Star Inc., Ashland, OR). Flow cytometry profiling and gating strategy is shown in Supplementary Fig. S1, S2, S4 and S5.

**Analysis of platelet uptake by macrophages in vitro**

Platelets isolated from WT or *Neu1*$^{Pf4\Delta Ex3}$ mice were labeled by PKH26 fluorescent dye (Sigma-Aldrich PKH26GL) following the manufacturer protocol. Then, the platelets in the concentration of $10^6$ cells/mL, premixed with anti-mouse CD41 monoclonal antibody at final concentration of 5 µg/ml, were added to the 6 well plates containing macrophages ($10^5$ cells/plate) isolated from WT or $Neu1^{Cx3cr1\Delta Ex3}$ mice and incubated at 37ºC or 4ºC for 30 min. After incubation, macrophages were washed three times with cold PBS to remove non-internalized platelets, detached by gentle scraping using a 1 mL syringe plunger, centrifuged at 300 × *g* for 5 min, resuspended in PBS, and fixed with 4% paraformaldehyde for 15 min at 4 °C. Cells were then analyzed by flow cytometry to quantify the percentage of PKH26-positive macrophages and the mean fluorescence intensity, reflecting the number of platelets engulfed per macrophage. In selected experiments, macrophages were pre-treated for 24 h or 30 min with NEU1 inhibitors including oseltamivir phosphate (OP), CG25901, CG33301, CG33300, and C9-BA-DANA. Synthetic inhibitors were prepared as described previously [28] and below. Compounds were originally reported as CG25900 (compound 11d), CG33300 (compound 17f), and C9-BA-DANA (compound 8) [28]. Compounds CG25901 and CG33301 are C1-methyl ester derivatives of their respective parents. $IC_{50}$ values were calculated from 3–5 independent biological experiments, each performed in technical triplicates. The experimental design and gating controls are shown in Fig. S1A.

**Passive murine ITP model.**

ITP in mice was induced as described by Katsman et al. [9]. Briefly, on the first day of the experiment, blood samples were collected from the submandibular vein to measure platelet counts by flow cytometry analysis as described above. Then, the mice were injected IP with escalating daily doses (68 μg/kg BW on the 1st day, 102 μg/kg dose on the 2d day) of mouse monoclonal anti-mouse CD41 antiplatelet antibody (clone MWReg30). The reticulated platelets were measured 6 h, 24 h and 48 h after the first injection of the antibodies by flow cytometric analysis. To assess the role of platelet surface desialylation by NEU1 in platelet clearance, ITP in platelet-specific conditional *Neu1* KO $Neu1^{Pf4\Delta Ex3}$ and control $Neu1^{loxPEx3}$ mice was also induced by anti-

GPIbα (anti-mouse CD42) antibodies that were reported to trigger NEU1 induction at the dose of 2 mg/kg BW. The reticulated platelets were measured 24 h after the injection as described above. To analyse the action of pharmacological neuraminidase inhibitors they were administered twice, 1 h before and 2 h after the injection of the platelet-depleting anti-CD41a antibodies or control mouse IgG in the dose of 68 μg/kg BW. The platelet concentration was measured by flow cytometry 6 h after the administration of the antibodies as described above. The experimental design and gating parameters are shown in Fig. S2.

**Neuraminidase 1 inhibitors**

Synthesis and characterization of the inhibitors used in this study (**CG33300** and **CG25900**) were previously reported in Guo et al. as **17f** and **11d**, respectively [10]. In the current study, we used the C1-methyl ester form of these compounds, which were prepared using a protocol identical to Guo et al., with the omission of the final deprotection of the methyl ester (NaOH, MeOH/$H_2O$).

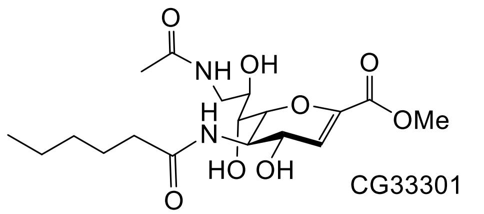


CG33301

**CG33301 $^1$H NMR (700 MHz, $CD_3OD$)** δ 5.94 (d, J = 2.5 Hz, 1H, H-3), 4.43 (dd, J = 8.7, 2.4 Hz, 1H, H-4), 4.18 (d, J = 10.6, 1H, H-6), 3.97 (dd, J = 10.8, 8.7 Hz, 1H, H-5), 3.93 (m, 1H, H-8), 3.78 (s, 3H, -$COCH_3$), 3.61 (dd, J = 14.0, 3.2 Hz, 1H, H-9), 3.41 (m, 1H, H7), 3.30 – 3.27 (m, 1H, H-9′), 2.28 (m, 2H, α-CH2), 1.97 (s, 3H, $COCH_3$), 1.65 (m, 2H, β-$CH_2$), 1.36 (m, 4H, γ-$CH_2$, δ-$CH_3$), 0.93 (t, J = 6.9 Hz, 3H, ε-$CH_3$).

**CG33301 $^{13}$C NMR (176 MHz, $CD_3OD$)** δ 178.11, 174.16, (2 × N−C=O), 164.31 (C-1), 145.18 (C-2), 113.64 (C-3), 78.03 (C-6), 71.68 (C-7), 69.99 (C-4), 67.78 (C-8), 52.76 (-$OCH_3$), 51.86 (C-5), 44.70 (C-9), 37.06 (C-α), 32.58 (C-β), 26.65 (C-γ), 23.41 (C-δ), 22.52 (-CO<u>C</u>$H_3$), 14.27 (C-ε).

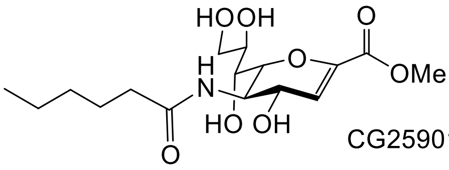


CG25901

**CG25901 $^1$H NMR (700 MHz, $CD_3OD$)** δ 5.95 (d, J = 2.3 Hz,1H, H-3), 4.43 (dd, J = 8.7, 2.3 Hz, 1H, H-4), 4.16 (dd, J = 10.8, 0.8Hz, 1H, H-6), 3.99 (dd, J = 10.8, 8.7 Hz, 1H, H-5), 3.91 (brs, 1H, H-8), 3.83 (dd, J = 11.4, 2.9 Hz, 1H, H-9), 3.78 (s, 3H, $COCH_3$) 3.63 (dd, J = 11.4, 5.5 Hz,1H, H-9′), 3.56 (dd, J = 9.3, 0.7 Hz, 1H, H-7), 2.29 (t, J = 7.5 Hz, 2H, α-$CH_2$), 1.69−1.62 (m, 2H, β-$CH_2$), 1.35 (m, 2 × 2H, γ-$CH_2$, δ-$CH_2$), 0.93 (t, J = 7.0 Hz, 3H, $CH_3$).

**CG25901 $^{13}$C NMR (176 MHz, $CD_3OD$)** δ 178.33 (N−C=O), 164.43 (C-1), 145.21 (C-2), 113.63 (C-3), 78.25 (C-6), 71.03 (C-7), 70.24 (C-4), 67.86 (C-8), 65.01 (C-9), 52.83 (-CO<u>C</u>$H_3$) 51.81(C-5), 37.04 (C-α), 32.54 (C- *β* ), 26.58 (C-γ), 23.43 (C-δ), 14.28 (C-ε).

The C1-methyl esters of DANA analogs are presumed to be cleaved by native esterases, similar to other prodrugs such as oseltamivir phosphate (an ethyl ester) [11]. Injection of WT mice (Male C57BL/6) with a C1-methyl ester (**CG14601**; IV 10 mg/mL in saline) and analysis of serum samples by HPLC showed a $T_{1/2}$ of 4.5 h; we also observed the appearance of the active (**CG14600**; $T_{max}$ 0.5 h, $T_{1/2}$ 1.8 h), consistent with conversion of the prodrug to the active form by native esterases.

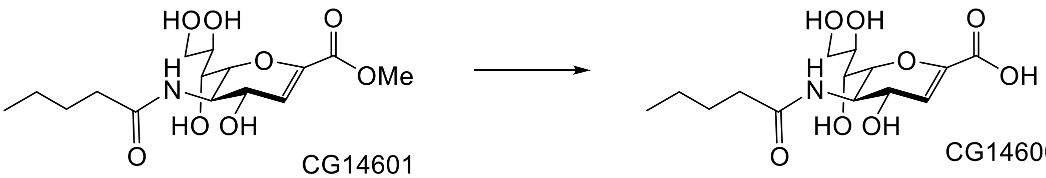


CG14601 CG14600

**Statistical analysis**

Statistical analyses were performed using Prism GraphPad 9.3.0. software (GraphPad Software San Diego, CA). The normality for all data was verified using the D'Agostino & Pearson omnibus normality test. Significance of the difference was determined using t-test (normal distribution) or Mann-Whitney test, when comparing two groups. One-way ANOVA or Nested ANOVA tests, followed by Tukey or Dunnett multiple comparison tests (normal distribution), or Kruskal-Wallis test, followed by Dunn multiple comparisons test, were used when comparing more than two groups. Two-way ANOVA followed by Tukey post hoc test was used for two-factor analysis. A *P*-value of 0.05 or less was considered significant.
